# Long term survival of *Dehalococcoides mccartyi* strains in mixed cultures under electron acceptor and ammonium limitation

**DOI:** 10.1101/2020.12.18.423565

**Authors:** Nadia Morson, Olivia Molenda, Katherine J. Picott, Ruth E. Richardson, Elizabeth A. Edwards

**Affiliations:** Department of Cell & Systems Biology, University of Toronto, Toronto, ON, Canada; Department of Chemical Engineering & Applied Chemistry, University of Toronto, Toronto, ON, Canada; School of Civil and Environmental Engineering, Cornell University, Ithaca, NY, United States of America

**Author notes:** Corresponding Author: Elizabeth A. Edwards, Department of Chemical Engineering and Applied Chemistry, University of Toronto, 200 College Street, Toronto, ON, Canada M5S 3E5.

**Keywords:** bioremediation, reductive dechlorination, quantitative PCR, *Dehalococcoides*, vinyl chloride

## Abstract

Few strains of *Dehalococcoides mccartyi* harbour and express the vinyl chloride reductase (VcrA) that catalyzes the dechlorination of vinyl chloride (VC), a carcinogenic soil and groundwater contaminant. The *vcrA* operon is found on a Genomic Island (GI) and therefore believed to participate in horizontal gene transfer. To try to induce horizontal gene transfer of the *vcrA*-GI, we blended two enrichment cultures in medium without ammonium while providing VC. We hypothesized that these conditions would select for a mutant strain of *D. mccartyi* that could both fix nitrogen and respire VC. However, after more than 4 years of incubation, we found no evidence for horizontal gene transfer of the *vcrA*-GI. Rather, we observed VC-dechlorinating activity attributed to the trichloroethene reductase TceA. Sequencing and protein modelling revealed a mutation in the predicted active site of TceA which may have influenced substrate specificity. We also identified two nitrogen-fixing *D. mccartyi* strains in the KB-1 culture. The presence of multiple strains of *D. mccartyi* with distinct phenotypes is a feature of natural environments and certain enrichment cultures (such as KB-1) and may enhance bioaugmentation success. The fact that multiple distinct strains persist in the culture for decades and that we could not induce horizontal gene transfer of the *vcrA*-GI suggests that it is not as mobile as predicted, or that mobility is restricted in ways yet to be discovered to specific sub-clades of *Dehalococcoides*.

**TOC Art:** 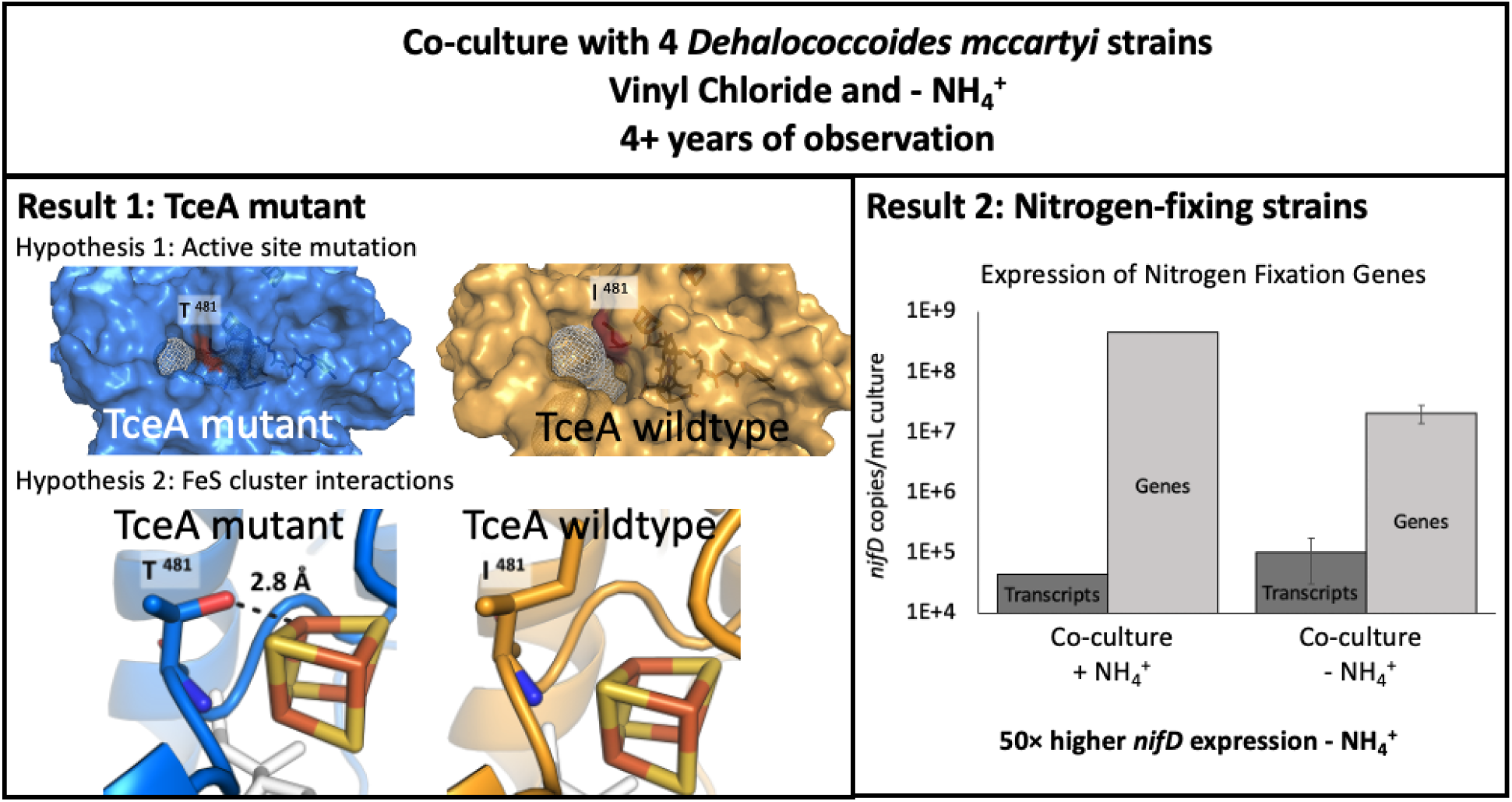

## Introduction

Chlorinated solvents are among the most prevalent and persistent soil and groundwater contaminants in industrialized countries (Moran, Zogorski and Squillace 2007). Tetra- or perchloroethene (PCE) and trichloroethene (TCE) contamination originates from the on-going use of dry-cleaning solvents and metal degreasing agents, respectively (Doherty 2000). These compounds, and their transformation intermediates such as vinyl chloride (VC), are known to have toxic or carcinogenic effects so their widespread soil and groundwater contamination poses a great concern to human health (Müller *et al.* 2004). In the late 1980s, following the discovery that microorganisms could completely dechlorinate these solvents under anaerobic conditions (Freedman and Gossett 1989), bioremediation and bioaugmentation emerged as highly successful treatment options for these problematic pollutants (Löffler and Edwards 2006). A microbial culture enriched from soil contaminated with chlorinated solvents in Southern Ontario, Canada, resulted in the KB-1™ mixed microbial consortium (Duhamel *et al.* 2002). KB-1™ has been used commercially for bioaugmentation for more than 20 years to remediate chlorinated compounds (Major *et al.* 2002; Löffler and Edwards 2006). In the KB-1™ culture, chlorinated ethenes, such as PCE and TCE, are sequentially dechlorinated via *cis*-dichloroethene (cDCE) and VC to ethene (Duhamel *et al.* 2002; Perez-de-Mora *et al.* 2017; Molenda *et al.* 2020). Reductive dechlorination in KB-1™ is primarily performed by multiple *Dehalococcoides mccartyi* strains in a growth-linked process called organohalide respiration (Duhamel *et al.* 2002; Perez-de-Mora *et al.* 2017; Molenda *et al.* 2020). Based on 16S rRNA sequence similarity, all *D. mccartyi* strains are categorized into three phylogenetic clades or sub-groups: Pinellas, Victoria, and Cornell. The Pinellas clade is represented by strain CBDB1, the Victoria clade is represented by strain VS, and the Cornell clade is represented by the type strain, strain 195 (Hendrickson *et al.* 2002; Löffler *et al.* 2013). Lab-grown KB-1 cultures enriched on different chlorinated substrates select for different strains of *D. mccartyi* that express different reductive dehalogenases (Perez-de-Mora *et al.* 2017). Reductive dehalogenases belong to a broad protein family (PF13486) and have been classified into Ortholog Groups (OGs) on the basis of >90% amino acid pairwise identity (Hug *et al.* 2013; Molenda *et al.* 2020). In the TCE-enriched KB-1 culture, strains belonging to the Pinellas clade are dominant, expressing the VC reductases VcrA (OG 8) and BvcA (OG 28) (Molenda *et al.* 2020). Recently, the TCE-enriched KB-1 culture was also found to contain low abundance Cornell clade strains, which express the TCE reductase TceA (OG 5) (Molenda *et al.* 2020). VC reductases are the most critical since VC is the most toxic dechlorination intermediate and following VC dechlorination, complete dechlorination is achieved.

Interestingly, the operon which encodes VcrA was found on a mobile genetic element, called the *vcrA*-Genomic Island (GI) (McMurdie *et al.* 2011). In the KB-1 TCE-enrichment culture, the *vcrA*-GI was identified in a circularized and extrachromosomal state within the cell which could theoretically be transferred between *D. mccartyi* strains through horizontal gene transfer (HGT) (Regeard *et al.* 2005; McMurdie *et al.* 2011; Molenda *et al.* 2019). However, there is no direct evidence of HGT of the *vcrA*-GI, and the mechanism of transfer between *D. mccartyi* remains unknown. To try to induce HGT of the *vcrA*-GI, we blended the KB-1 TCE-enrichment culture with another mixed culture, called Donna II (Fennell, Gossett and Zinder 1997). We called the blend of the two enrichment cultures DKB (Donna + KB-1). The KB-1 TCE-enriched consortium is the ideal *vcrA*-GI donor because the *vcrA*-containing *D. mccartyi* strain is highly abundant in this culture. The Donna II culture is a mixed microbial consortium that contains only one *D. mccartyi* strain, strain 195, that dechlorinates PCE to VC via organohalide respiration utilizing the PCE dehalogenase PceA (OG 30) and TceA (Fennell, Gossett and Zinder 1997). Subsequently, VC is dechlorinated to ethene slowly via co-metabolism since *D. mccartyi* strain 195 does not contain the *vcrA*-GI or any other VC reductases and therefore cannot grow on VC (Regeard *et al.* 2005). Of particular interest for experimental design, *D. mccartyi* strain 195 is capable of nitrogen fixation in the absence of available nitrogen sources via the *nif* operon (Lee *et al.* 2009). Whereas the *vcrA*-GI-containing *D. mccartyi* strain from TCE-enriched KB-1 does not have a nitrogen fixation operon and thus an incapability for nitrogen fixation. In a theoretical HGT event, the KB-1 TCE-enrichment culture would contain the donor strain of the *vcrA*-GI and *D. mccartyi* strain 195 in Donna II would be the recipient strain. By applying the selective pressure of providing only VC as the sole energy source (electron acceptor) in medium without ammonium, we thought that a hybrid *D. mccartyi* strain which could fix nitrogen and respire VC would emerge.

After 4 years of observation, we found no evidence to support horizontal gene transfer of the *vcrA*-GI. Instead, in two of three replicates of the DKB culture, we observed VC dechlorination activity even though *vcrA* gene copies were low and *tceA* and *D. mccartyi* 16S rRNA copies were high. We then sequenced and modelled the TceA of the DKB culture and found a mutation in the predicted active site that is hypothesized to influence substrate specificity. Additionally, over the course of the study, two previously unknown Cornell strains of *D. mccartyi* were identified in KB-1 that contain nitrogen-fixing genes. We determined that these genes were expressed and active at low ammonium concentrations, a beneficial feature of the KB-1 culture for bioremediation.

## Materials and Methods

### Enrichment cultures

The KB-1 culture originated from microcosms prepared with aquifer materials from a TCE-contaminated site in southern Ontario in 1996 (Duhamel *et al.* 2002). The KB-1 TCE-enrichment culture is maintained biweekly with 0.76 mM TCE as the electron acceptor and 5 × electron equivalents of methanol (MeOH) as the electron donor, referred to as KB-1/TCE-MeOH, as previously described (Duhamel, Mo and Edwards 2004; Duhamel and Edwards 2006, 2007). The Donna II culture originated from an enrichment culture seeded with digester sludge from a wastewater treatment plant in Ithaca, NY, USA. The Donna II culture was maintained batch-style at Cornell University with 110 μM of PCE as the electron acceptor and butyrate as the electron donor, as previously described (Smatlak, Gossett and Zinder 1996; Fennell, Gossett and Zinder 1997). The “DKB” culture was formed at Cornell University in 2012 when 1000 mL of KB-1/TCE-MeOH and 700 mL of Donna II were combined. The DKB culture was maintained with PCE and butyrate, similar to the Donna II culture. In 2015, the DKB culture was shipped to the University of Toronto and was subsequently used as inoculum to create DKB sub-cultures grown under different conditions of varying selective pressures, as described below.

### Experimental Set-up and Monitoring

DKB sub-cultures were fed either PCE or VC as electron acceptor, and butyrate as electron donor. Additionally, DKB sub-cultures were either cultured in medium with ammonium (▲), or without ammonium (Δ) requiring nitrogen fixation. Therefore, four experimental conditions were prepared: (i) DKB PCE ▲, (ii) DKB PCE Δ, (iii) DKB VC ▲, and (iv) DKB VC Δ (Figure S1). Additionally, a KB-1 control culture, KB-1/TCE-MeOH, was used as inoculum for sub-culture triplicates into medium without ammonium and amended with VC, referred to as KB-1 VC Δ; this condition was expected to be a negative control for KB-1 growth under VC-degrading, ammonium-limiting conditions (Figure S1). To create each sub-culture, each 2 mL sample of culture (KB-1/TCE-MeOH or DKB) was centrifuged at 13,000 × g for 15 minutes at room temperature, the supernatant was discarded, and the pellet was resuspended by flicking and pipetting in 2 mL of anaerobic, autoclaved distilled H_2_O. This process was repeated three times, to wash ammonium from the pellet. Following, the 2 mL volume of washed inoculum was transferred into 198 mL of anaerobic mineral medium (Duhamel and Edwards 2007) with (10 mM) or without NH_4_Cl in 250 mL serum bottles sealed with butyl rubber stoppers (Geo-Microbial Technologies Inc.). An additional 500 μL of vitamin stock (Edwards and Grbic-Galic 1994) was added to each bottle during set-up, and 50 μL was added for each 20 mL of medium added during medium amendments. The culture was purged with N_2_:CO_2_ (80:20) gas, and the headspace was over-pressurized with N_2_:CO_2_ for 3 seconds. The DKB sub-cultures were amended periodically with 310 μelectron equivalents (μeeq) of gaseous VC (5 mL) or neat PCE (4 μL) as electron acceptor, and 4 × eeq sodium butyrate stock as electron donor. Electron equivalents were used to calculate the mass of electron donor or acceptor fed to the cultures as a way to establish a ratio of electrons required for organohalide respiration given that each hydrogenolysis reaction requires 2 electrons for the removal of a chlorine atom. Therefore, the KB-1/TCE-MeOH sub-cultures, KB-1 VC Δ, were maintained with 310 μeeq VC as electron acceptor, and 5 × eeq of methanol and ethanol mixture (50:50 on an eeq basis) as electron donor. Cumulative electron acceptor consumed was monitored over time. Acceptor and donor were re-amended as needed when depleted in each sub-culture, and medium was amended following large volume sampling for nucleic acid extraction in batch style maintenance. Triplicate bottles were prepared for each of the four test conditions. In 2016, to reduce number of bottles monitored, only one of the triplicate cultures from the DKB sub-cultures with ammonium was maintained; prior to this time, these triplicates were behaving similarly. Therefore, from 2016 onward, 8 DKB sub-cultures and three KB-1 VC Δ cultures were maintained and analyzed.

### DNA Extraction

Samples were taken from KB-1/TCE-MeOH, Donna II, DKB, and DKB sub-cultures at various times throughout the experiment for DNA extraction. For each DNA sample, a volume of 20 mL of liquid culture was removed anaerobically. The 20 mL aliquot was centrifuged in a sealed Falcon conical centrifuge tube at 6870 × g for 20 minutes at room temperature, using a swinging-bucket rotor. Aerobically, the supernatant was decanted, and the pellet was resuspended immediately using 60 μL of Solution C1 and 500 μL of liquid from the PowerBead tube, from the DNeasy PowerSoil Kit (Qiagen). All subsequent kit instructions were followed. Final DNA elution volume was 50 μL of Solution C6 (elution buffer). DNA was quantified by NanoDrop or using the High-Sensitivity DNA kit with the Qubit® 3.0 Fluorometer (ThermoFisher Scientific). All DNA extracts were stored at −80°C.

### RNA Extraction and cDNA synthesis

RNA was extracted from 10 mL of the KB-1 VC Δ cultures and the KB-1/TCE-MeOH culture. Cells were pelleted by centrifuging at 5000 × g at 4°C for 15 minutes. Pellets were then resuspended in 500 μL of supernatant and stabilized with 1 mL of RNAprotect Bacteria Reagent (Qiagen). The RNeasy® Protect Bacteria Mini Kit (Qiagen) was used to extract RNA from the cells by physical disruption via bead-beating. The kit procedure was followed, and RNA was eluted in 30 μL of RNase-free water. Following, RNA was DNase-treated using 10 μL of DNase A and 70 μL of DNase buffer for 30 seconds. To purify, the RNA was cleaned up using the RNeasy® MinElute® Cleanup kit (Qiagen) and eluted in 15 μL of RNase-free water. Following, cDNA was synthesized from the RNA using the Invitrogen Superscript® VILO cDNA synthesis kit (Invitrogen), and procedure was followed according to kit instructions. cDNA was quantified using the High-Sensitivity DNA kit with the Qubit® 3.0 Fluorometer. All cDNA extracts were stored at −80°C.

### *D. mccartyi* strain biomarker selection for Quantitative PCR of 16S rRNA and functional genes

To track the growth of the different *D. mccartyi* strains in the DKB sub-cultures, we designed strain-specific biomarkers for quantitative PCR (qPCR). The abundance of all *D. mccartyi* strains was quantified by the 16S rRNA gene, using primers Dhc1f and Dhc264r (Hendrickson *et al.* 2002), since there is a single 16S rRNA gene copy per genome. To quantify the *vcrA*-containing strain from KB-1/TCE-MeOH and all extrachromosomal *vcrA*-GIs, we used primers vcrA670f and vcrA440r targeting the *vcrA* gene (Molenda, Quaile and Edwards 2016). The tceA500f and tceA795r (Fung *et al.* 2007) primers were used for tracking *tceA*-containing *D. mccartyi* strains in KB-1/TCE-MeOH and *D. mccartyi* strain 195. To quantify strain 195, we designed primers for a unique *rdhA* (DET_RS00960), that is truncated (truncated *rdhA*, “t*rdhA”*) and predicted to encode a non-functional reductive dehalogenase. It is only 1200 bp long compared to functional *rdhA* with lengths of approximately 1500 bp. As well, this gene contains two iron-sulfur cluster binding domains (CX_2_CX_2_CX_3_CP)_2_ but does not contain commonly conserved motifs: a twin-arginine TAT membrane export sequence (RRXFXK) nor a cobalamin binding domain. Primers for t*rdhA* were designed for the *D. mccartyi* strain 195 genome, using Primer2 (Untergasser *et al.* 2012) in Geneious 8.1.9 (Kearse *et al.* 2012). To track nitrogen fixation (*nif)* genes, we quantified the *nifD* gene which encodes the nitrogenase molybdenum-iron protein ⍰-chain. The *nifD* primers used in this study were used to characterize nitrogen fixation in *D. mccartyi* strain 195 (Lee *et al.* 2009). These *nifD* primers were also used for transcription analysis of nitrogen fixation genes in KB-1 VC Δ and KB-1/TCE-MeOH, by reverse transcription (RT)-qPCR. Lastly, we went back to archived DNA samples to quantify the vinyl chloride reductase, *bvcA* (Figure S2), using primers bvcA318f and bvcA555r (Waller *et al.* 2012). More information on primers can be found in Supplementary Information (Table S1).

As a qPCR standard for absolute quantification, a plasmid with concatenated sequences corresponding to the *D. mccartyi* 16S rRNA gene, *vcrA*, *tceA,* and *bvcA* was used, as previously described (Molenda *et al.* 2019). This concatenated plasmid allowed us to calculate accurate ratios of these *rdhA* genes to 16S rRNA gene copies. For *nifD* and t*rdhA* qPCR standards, regions were PCR amplified, purified and cloned into *Escherichia coli* (TOPO™ TA Cloning™ Kit for Sequencing, Invitrogen) (Additional methods in Supplemental Information).

All qPCRs were prepared in a UV PCR cabinet and each qPCR was run in duplicate or triplicate, using a CFX96 real-time PCR detection system. Each 20 μL qPCR reaction was prepared in UV-treated UltraPure nuclease-free water containing 10 μL of EvaGreen® Supermix (Bio-Rad Laboratories, Hercules, CA), 0.5 μL of each primer (forward and reverse, from 10 μM stock solutions), and 2 μL of DNA template or standard plasmid dilution series, from 10^1^ to 10^7^ copies of plasmid per μL. Thermocycler program: initial denaturation at 95°C for 2 min, followed by 40 cycles of denaturation at 98°C for 5 s, varied annealing temperatures (Table S1), followed by extension for 10 s at 72°C. All qPCR standards and quality metrics (efficiency, standard curve details, etc.) can be found in Supplementary Information (Table S2), with calculations of absolute gene abundances per mL of culture (Table S3). The quantification limit (corresponding to lowest calibration standard) was ~1×10^3^ copies/mL, but values as low as 1×10^1^ copies/mL could be detected. To compare *nifD* expression among different experimental conditions, the absolute *nifD* gene abundances of genomic DNA and cDNA were determined. Using this data, we calculated *nifD* transcript per gene (TPG) ratios, as the ratio of transcript copies per mL of culture to gene copies per mL of culture.

### Clone Library Preparation and Sequencing of *tceA*

A clone library of *tceA* was generated from one of the DKB VC Δ cultures. As control, a *tceA* clone library was generated in parallel using the KB-1/TCE-MeOH culture. The two *tceA* clone libraries were generated using the TOPO™ XL-2 Complete PCR Cloning Kit (Invitrogen). PCR primers were designed to amplify the 2200 bp *tceAB* region (Table S1). Blunt end *tceA* PCR products were produced using the Platinum SuperFi polymerase, bands were extracted from an agarose gel, and cloned into One Shot™ OmniMAX™ 2 T1R Chemically Competent *E. coli* cells. 50 μL or 100 μL of transformed *E. coli* were spread on 50 μg/mL kanamycin LB plates, with 40 μL of 40 mg/mL X-gal in dimethylformamide (DMF) solution and 40 μL of filter-sterilized 0.1 M IPTG for blue-white selection. Following, 10 colonies were selected each from the DKB sub-culture and from the KB-1/TCE-MeOH culture and transferred cultures onto a patch plate for further analysis. From the patch plate, colonies were incubated in to 10 mL of 50 μg/mL kanamycin LB broth, and plasmids were extracted using a QIAprep Spin Miniprep Kit (Qiagen). Each plasmid was PCR amplified using the T3 and T7 cloning primers to confirm successful transformation. For the first round of Sanger sequencing, plasmids were sequenced using the T3 and T7 cloning primers, at the SickKids Center for Applied Genomics (TCAG) sequencing/synthesis facility (Toronto, Canada). Ten sequences each from the forward and reverse were then aligned, and two consensus sequences were generated at a 99% nucleotide identity threshold. Using the consensus sequences, primers were designed (Table S1) for the remaining 845 bp. For the second round of sequencing, the final 99% nucleotide identical consensus sequences were generated by aligning these 20 sequences. The cloned *tceA* sequences from this study, the *tceA* published on NCBI (AAW39060), and the *tceA* from the Donna II metagenome (IMG-M taxon ID: 2032320001) were aligned using the MUSCLE aligner (Edgar 2004) in Geneious 8.1.9.

### TceA protein modelling

Protein models were independently produced for TceA from the *Dehalococcoides mccartyi* 195 isolate and the TceA from one of the DKB VC Δ cultures. Each sequence was first trimmed to remove the TAT signal peptide sequence predicted by SignalP-5.0 (Almagro Armenteros *et al.* 2019). The models were produced from four different predictive modelling servers, AlphaFold2, Robetta, I-TASSER, and Phyre2, and assessed based on quality scores determined by the MolProbity assessment tool (Chen *et al.* 2010; Roy, Kucukural and Zhang 2010; Song *et al.* 2013; Kelley *et al.* 2015; Studer *et al.* 2020; Jumper *et al.* 2021). The AlphaFold2 and Robetta models had the highest quality scores and were used for further analysis; however, all models were used to validate the predicted spatial locations of the mutated residues. The cobalamin cofactor was modelled into the active site using AutoDock Vina which was consistent with the docking position of known crystal structures (Trott and Olson 2009). The iron-sulfur clusters were superimposed from the *Sulfurospirillum multivorans* PceA crystal structure (PDB ID: 4UR2) (Bommer *et al.* 2014). The substrate access channels were predicted using the CAVER 3.0 plugin in PyMol v2.3.4, only the channels predicted on the catalytic face of the cobalamin cofactor were kept (Chovancova *et al.* 2012). Models were imaged and the molecular interactions were predicted using PyMol v2.3.4 (The PyMOL Molecular Graphics System, Version 2.3.4 Schrödinger).

### Nitrogenase operon gene and protein alignments

The *D. mccartyi* strain 195 nitrogen fixation operon (DET_RS05950-5990), encoding the nitrogenase enzyme, was aligned with the nitrogenase operons of the two KB-1/TCE-MeOH strains (KBTCE2: B1773_05150-05185, KBTCE3: B1774_04800-04835). An alignment of these operons was performed using the built-in MUSCLE aligner (Edgar 2004) in Geneious 8.1.9.

## Results and Discussion

### Long term growth of DKB sub-cultures under varying degrees of selective pressure

To analyze the effects of experimental conditions on growth and dechlorination, we monitored the cumulative electron acceptor consumed over 4 years (Table S3). Overall, cultures grown without ammonium (Δ) did not consume as much electron acceptor as cultures with ammonium (▴) in each series (Figure 1A). The DKB PCE ▴ culture consumed twice as much PCE as the DKB PCE ▴ cultures, while the DKB VC ▴ culture consumed nearly six times more VC than the DKB VC Δ cultures over the same time. The DKB VC Δ cultures, grown under the highest selective pressure conditions of VC without ammonium, dechlorinated the least amount of electron acceptor of all DKB sub-cultures (Figure 1A). These findings confirmed that the selective pressures on the DKB sub-cultures impacted the metabolism of *D. mccartyi* to varying degrees.

**Figure 1:**
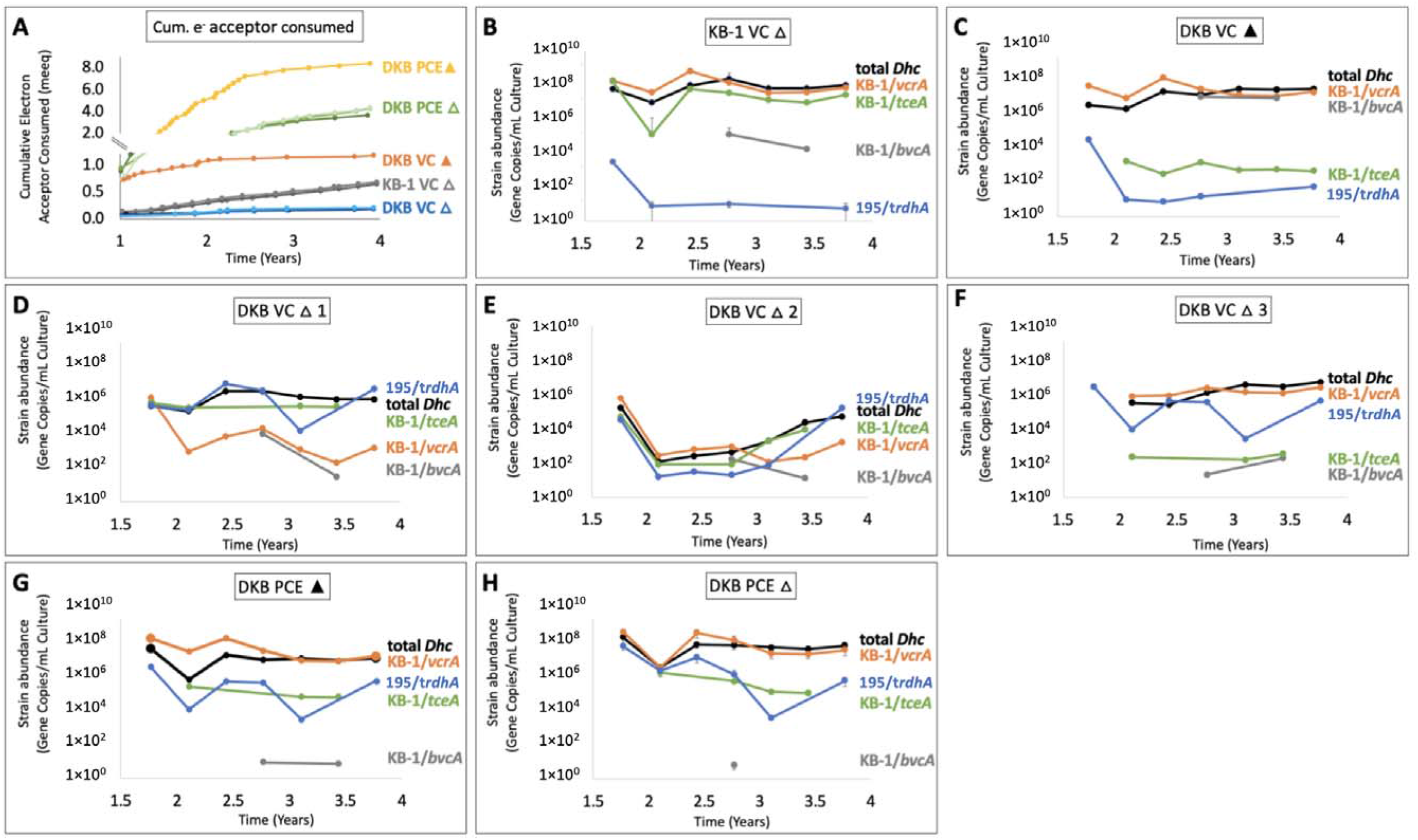
Cumulative electron acceptor consumed (meeq) and strain-specific biomarker tracking (gene copies/mL culture). (A) Cumulative electron acceptor consumed, given that 8 eeq are required per mol of PCE consumed, and 2 eeq per mol VC consumed. (B) *D. mccartyi* abundance, by 16S rRNA gene copies per mL of culture. (C) *vcrA* copies per mL of culture, quantifying both genomicic and extrachromosomal *vcrA*-GI. (D) *D. mccartyi* strain 195-specific biomarker for a “truncated” *rdhA* (t*rdhA*). (E) *tceA* and (F) *nifD* gene copies per mL of culture. Closed triangles represent ammonium provided in the medium (▴), and open triangles represent ammonium not provided (Δ) in growth medium. The quantification limit was 1×10^3^ copies/mL; values below this are unreliable. Error bars represent the standard deviation (n=3), otherwise unaveraged triplicates are numbered (1,2,3) or a single bottle if not numbered (n=1).

In each DKB and KB-1 culture, we tracked the abundance of key genes of *D. mccartyi* strains and related that to strain abundance (Figure 1B–H). The *D. mccartyi*-specific 16S rRNA gene target tracked the abundance of all *D. mccartyi* strains (“total *Dhc*”). The *vcrA* gene target provided an indication of *D. mccartyi* strain KBTCE1 from KB-1 that contained the *vcrA*-GI (“KB-1/*vcrA*”) and captured any extrachromosomal copies of the *vcrA*-GI. Similarly, the *bvcA* gene target measured the *bvcA*-containing strain from KB-1 (“KB-1/*bvcA*”). *D. mccartyi* strain 195 and two KB-1 strains, KBTCE2 and KBTCE3, contained both *tceA* and *nifD* gene targets, so the strain 195 biomarker (truncated *rdhA*, or t*rdhA*) was used to distinguish between *D. mccartyi* from KB-1 (“KB-1/*tceA*”) and strain 195 (195/t*rdhA*”).

Trends in the abundance of key strain biomarkers helped decipher what occurred in the DKB (blend of KB-1 and Donna II) and KB-1 control cultures over nearly 4 years of incubation. The KB-1/*vcrA* strain was highly abundant (10^7^ to 10^9^ copies/mL of culture) in all cultures except the DKB VC ε (no ammonium) replicate cultures (Figure 1D–F). In DKB VC Δ1 and Δ2 cultures (Figure 1D–E), the KB-1/*vcrA* strain decreased in abundance over time, whereas in the DKB VC Δ3 culture, KB-1/*vcrA* abundance was maintained (Figure 1F). Conversely, an increase in the 195/t*rdhA* strain was observed in DKB VC Δ1 and Δ2 cultures, whereas it was below or near the quantification limit of 10^3^ copies/mL in the DKB VC Δ3 culture throughout the experiment (Figure 1D–F). All together, these results suggest that the dominant strain in DKB VC Δ1 and Δ2 cultures was *D. mccartyi* strain 195 from the Donna II culture, and the dominant strain in the DKB VC Δ3 culture was *D. mccartyi* strain KBTCE1 from the KB-1 culture (Figure S3). An interesting and complementary result is that the non-dominant *D. mccartyi* strains, such as KB-1/*bvcA* and KB-1/*tceA*, persisted in low abundances throughout the 4 years of incubation. If the *vcrA* island were to be transferred to strain 195, the “hybrid” *D. mccartyi* strain would have both *vcrA* and t*rdhA* and thus would appear in Figures 1D–F as an equal abundance of KB-1/*vcrA* and 195/t*rdhA,* yet this result was never observed. Therefore, these results refute the existence of a predicted hybrid strain that acquired the *vcrA*-GI.

However, early in the experiment, we noticed that KB-1 VC Δ cultures were able to dechlorinate VC even without ammonium provided in the growth medium (Figure 1A, Table S4). This condition was intended to be a control where dechlorination and growth were not anticipated. As dechlorination continued, it was suspected that the KB-1 VC Δ culture may contain nitrogen-fixing bacteria, later justified by the high abundance of *D. mccartyi nifD* copies/mL of culture (Figure 1F). We therefore analyzed nitrogen fixation activity in the KB-1/TCE-MeOH and KB-1 VC Δ cultures.

### Complete nitrogenase operons identified in two KB-1 strains

In 2012, when this experiment was conceived, the only *D. mccartyi* strain that was known to fix nitrogen was the *D. mccartyi* strain 195 isolate (Lee *et al.* 2009). After the DKB experiment had begun, metagenomic sequencing of the KB-1/TCE-MeOH culture revealed the presence of more than three strains of *D. mccartyi* (Molenda *et al.* 2019). The *vcrA*-GI containing strain, KBTCE1, was the most abundant strain and belonged to the Pinellas clade. However, the other two strains, KBTCE2 and KBTCE3, were both *tceA-*containing Cornell strains that were not previously detected in the culture. Genome analysis revealed that strains KBTCE2 and KTBCE3 contained complete nitrogenase operons, encoded by *nif* genes, and were therefore predicted to fix nitrogen. The nitrogenase operons in strains KBTCE2 and KBTCE3 share 99.0% nucleotide pairwise identity with the Cornell strain 195 isolate nitrogenase operon (DET_RS05950-RS05990) (Figure 2A, Table S8). The KBTCE2 and KBTCE3 nitrogenase operons were almost identical to each other, with one single nucleotide polymorphism (SNP) in the *nifK* gene (Figure 2A), which is the ⍰-subunit of the iron-molybdenum protein in the nitrogenase complex (Raymond *et al.* 2004).

**Figure 2:**
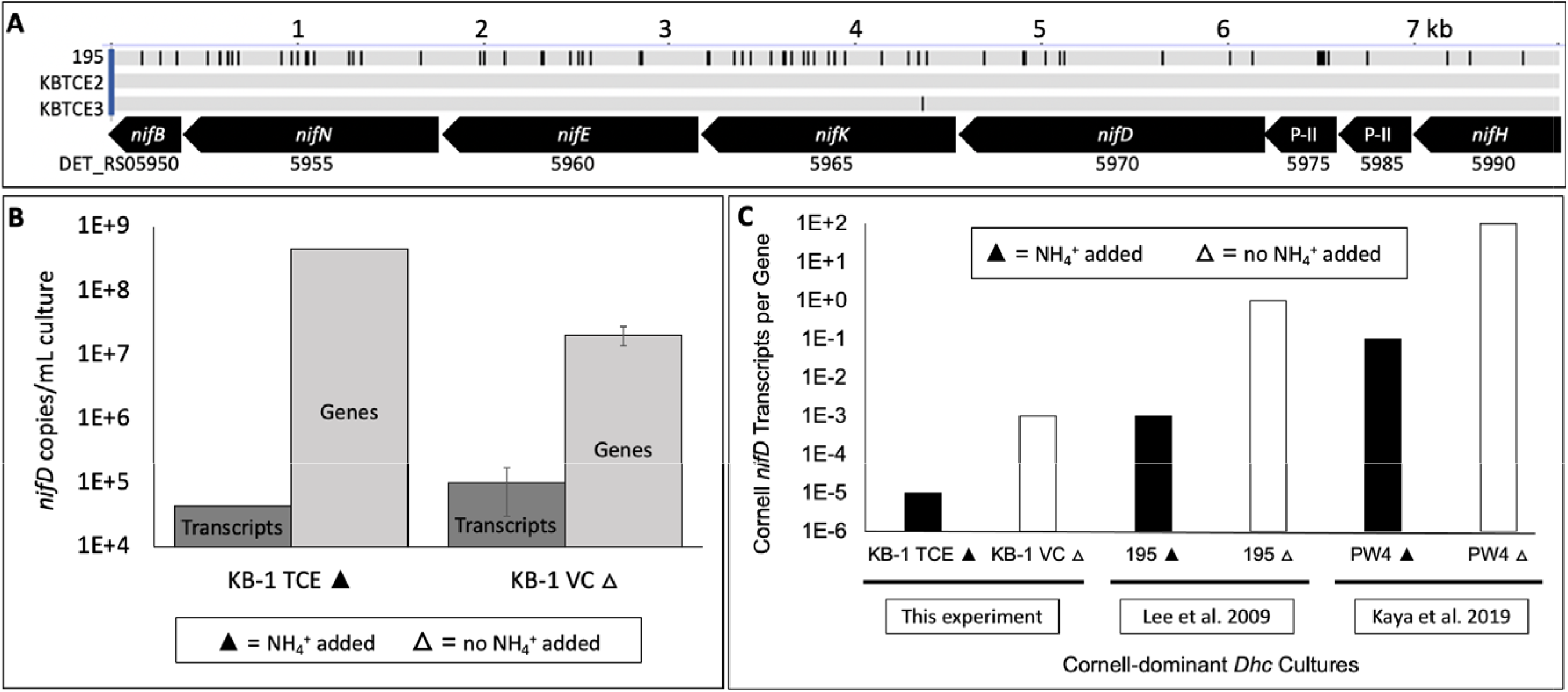
Nitrogen fixation in *D. mccartyi* (*Dhc*) in KB-1. (A) Alignment of nitrogenase genes (*nif*) in *D. mccartyi* strains 195 (DET_RS05950-5990), KBTCE2 and KBTCE3, where each black bar represents a misaligned nucleotide position. The top scale indicates nucleotide length in kilobase pairs (kb). (B) *nifD* transcripts and gene copies per mL of KB-1 grown with (▲) and without (ε) ammonium provided. Error bars represent standard deviation (n=3). (C) *nifD* transcripts per gene (TPG) in KB-1, compared to approximate TPG ratios from Cornell-dominant *Dhc* cultures: pure strain 195 culture (Lee *et al.* 2009) and mixed PW4 culture (Kaya *et al.* 2019), grown with (▲) and without (Δ) ammonium.

To determine if KBTCE2 and KBTCE3 strains could utilize the *nif* operon to fix nitrogen, we quantified the *nifD* gene copies and transcripts in the KB-1 VC Δ cultures and the KB-1/TCE-MeOH culture, referred to here as KB-1 TCE ▴. In the KB-1 VC Δ cultures, the absolute abundance of *nifD* transcripts was 2.3 (± 1.7) times more than the KB-1 TCE ▴ culture, even though the absolute abundance of the *nifD* gene was 24 (± 9.6) times less (Figure 2B).

Therefore, *D. mccartyi* strains KBTCE2 and KBTCE3 in the KB-1 VC Δ cultures were actively transcribing the *nif* operon indicative of nitrogen fixation. However, the KBTCE2 and KBTCE3 strains are *tceA*-containing strains and do not contain *vcrA*, so these strains are not able to obtain energy from VC in these experimental conditions. This is different from the KB-1 TCE ▴ culture in which strains KBTCE2 and KBTCE3 can obtain energy through the dechlorination of TCE. Therefore, it is a conundrum as to how these strains obtain energy when grown in the presence of VC as the sole electron acceptor in the KB-1 VC ε cultures, especially since nitrogen fixation is an energetically expensive process (Leigh and Dodsworth 2007). This finding might suggest syntrophic relationships between *D. mccartyi* strains, in the exchange of energy for available nitrogen sources. Alternatively, this activity may point towards the function of the constitutively expressed reductive dehalogenase (OG 15) observed under “starvation” conditions in *D. mccartyi* strain 195 (DET_RS07915) (Johnson *et al.* 2008; Rahm and Richardson 2008) and in KB-1 (DQ177510) (Waller *et al.* 2012; Liang *et al.* 2015). It may be that RdhA from OG 15 can sustain the population with an unknown electron acceptor in what is perceived as “starvation”. Further work is required to determine if a reductive dehalogenase from OG 15 is expressed in the KB-1 VC Δ cultures and its role in providing energy to *D. mccartyi* strains.

The *nifD* transcripts-per-gene (TPG) ratio was calculated for the KB-1 VC Δ cultures and for KB-1 TCE ▴ (Figure 2C). The TPG ratios of KB-1 cultures were comparable to approximate TPG ratios of the *D. mccartyi* strain 195 isolate (Lee *et al.* 2009) and the PW4 groundwater aquifer-derived enrichment culture dominated by Cornell strains (Kaya *et al.* 2019) (Figure 2C, Table S6). Therefore, we identified two more strains of *D. mccartyi* that can fix nitrogen, both originating from the KB-1/TCE-MeOH culture. *D. mccartyi* strains KBTCE2, KBTCE3, and 195 are the only strains currently known to fix nitrogen, all belonging to the Cornell clade (Molenda *et al.* 2020). Therefore, the nitrogen fixation characteristic of *D. mccartyi* appears to be specific to the Cornell clade.

### *D. mccartyi* strain 195 TceA predicted VC-dechlorinating activity

*D. mccartyi* strain 195 in the Donna II culture contains *tceA* and *D. mccartyi* strains KBTCE2 and KBTCE3 in the KB-1/TCE-MeOH also contain *tceA* (Molenda *et al.* 2020). The *tceA* gene encodes the TCE reductive dehalogenase, TceA, which catalyzes the reductive dechlorination of TCE to cDCE and cDCE to VC (Tang *et al.* 2013). TceA can also dechlorinate 1,2-dichloroethane (1,2-DCA) to ethene and trace amounts of VC (Duhamel and Edwards 2007). However, it was recently discovered that TceA can dechlorinate VC to ethene coupled to organohalide respiration in the presence of sufficient vitamin B_12_ (Yan *et al.* 2021). Ethene formation occurred when vitamin B_12_ concentrations were 10 μg/L or greater and dechlorination rates were positively correlated to B_12_ concentrations (Yan *et al.* 2021). In the DKB experiment, vitamin B_12_, in the form of cyanocobalamin, was maintained at a concentration of 6 μg/L in the culture medium, which would suggest that VC would not be consumed. However, our data revealed slow VC dechlorination where ethene was formed at a rate of 0.88 (± 0.58) μmol/day in the DKB VC Δ1 culture (Figure S4) which was comparable to the rate of Cl^-^ released when vitamin B_12_ concentrations were 10 μg/L (Yan *et al.* 2021). Furthermore, the observed increase in *tceA* and *D. mccartyi* with the simultaneous decrease in *vcrA* (Figure 1D–F) suggested that *D. mccartyi* were able to sustain growth on VC dechlorination with *tceA* with 6 μg/L of vitamin B_12_ (Figure S4).

To determine if the active TceA was from the Donna II or KB-1 culture, we generated a *tceA* clone library from the DKB VC Δ1 culture (Table S7) and the KB-1/TCE-MeOH culture as a control. The cloned *tceA* sequence from the KB-1/TCE-MeOH culture was identical (100% nucleotide pairwise identity) to the *tceA* in published KBTCE2 and KBTC3 genomes (Figure S5A). The cloned *tceA* sequence from the DKB VC Δ1 culture was 100% identical to the *tceA* in *D. mccartyi* strain 195 from the Donna II metagenome sequenced in year 2010 (Figure S5B), but not the original sequence of isolated strain 195, sequenced in 2005. Therefore, the observed VC-dechlorinating activity was not the result of a mutation during this experiment. TceA from the Donna II metagenome was aligned to TceA from the isolated *D. mccartyi* strain 195 and other TceA from OG 5 (Figure S6). In this alignment, two non-synonymous mutations were identified between the strain 195 TceA and Donna II TceA: V261A and I481T. Interestingly, the I481T mutation was also observed in the recently sequenced TceA of strain FL2, which was shown to have higher rates of VC respiratory activity compared to strain 195, even at high vitamin B_12_ concentrations (Yan *et al.* 2021).

To explore whether these two mutations could have an impact on the substrate specificity and activity of the protein, we generated models of the TceA structure using the sequence from *D. mccartyi* strain 195 isolate and from the Donna II metagenome. Models were generated independently through several prediction servers to increase confidence in the predicted locations of each mutation (Table S9), with the highest scoring models produced by the Robetta and AlphaFold2 servers (Song *et al.* 2013; Jumper *et al.* 2021). In each model the I481T mutation was consistently near the active site in both the *D. mccartyi* strain 195 isolate and the Donna II strain 195 TceA models (Figure 3B). Whereas the location of the V261A mutation had some variability but was always predicted to be removed from the active site. It is unlikely that the V261A mutation plays a role in altering the enzymatic activity (Figure 3A). Visualization of the predicted I481T location led to two potential hypotheses that could contribute to enhanced VC respiration: alteration of the substrate access channel, and hydrogen bonding between the residue and an [4Fe-4S] cluster to shift its reduction potential. These are described below.

**Figure 3:**
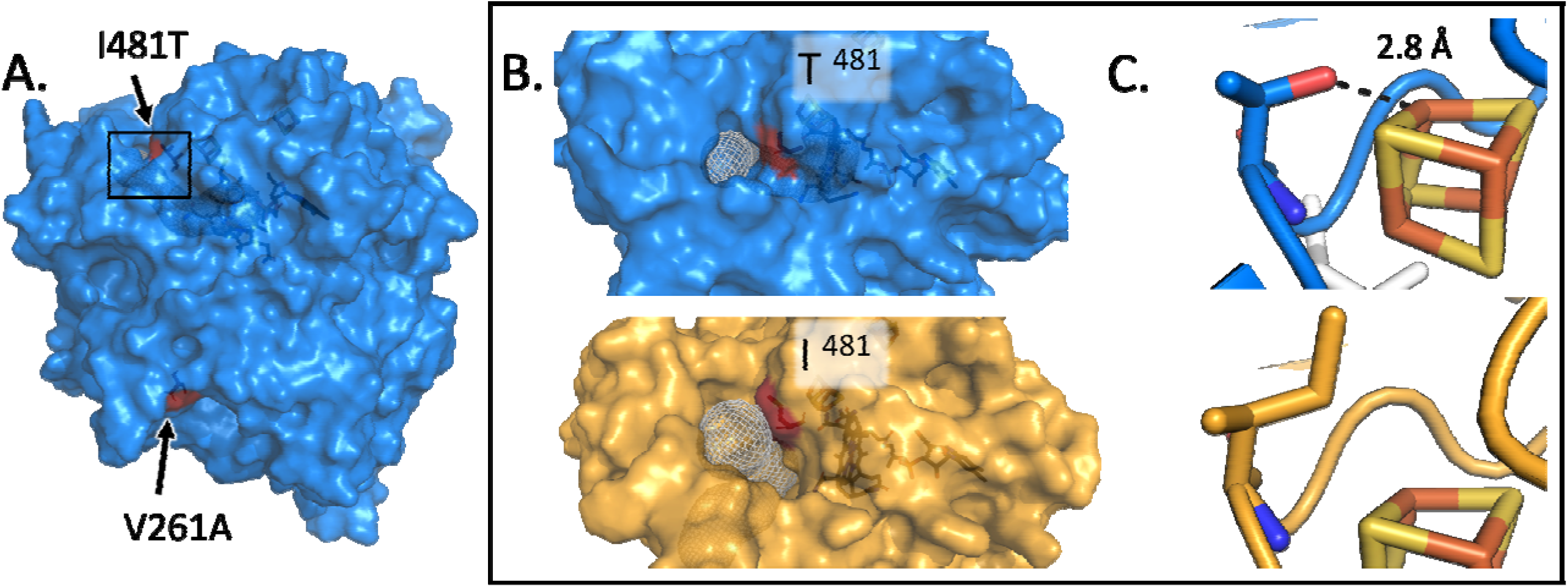
Protein models of the Donna II metagenome *D. mccartyi* strain 195 TceA (blue) and *D. mccartyi* strain 195 isolate TceA (yellow) with mutation sites highlighted in red. (A) The Donna II TceA mutant structure shows spatial locations of V261A and I481T mutations. (B) View of the substrate access channel (depicted in white mesh) with the I481T mutation (red) in the Donna II TceA and strain 195 isolate TceA. (C) Close up of the two residues next to a superimposed [4Fe-4S] cluster with the measured distance (2.8 Å) indicating the distance of the T481 hydroxyl group to the cluster. Models were produced using the Robetta and AlphaFold2 online prediction servers, [4Fe-4S] clusters were superimposed from the PceA crystal structure (PDB ID: 4UR2) (Song *et al.* 2013).

The crystal structure of PceA, a PCE reductive dehalogenase, from the organohalide-respiring bacterium *Sulfurospirillum multivorans,* depicts a narrow substrate channel leading to the buried hydrophobic binding pocket (Bommer et al., 2014). To visualize where the substrate may enter the active site in our models, we predicted the structure all of the tunnels with access to the catalytic face of the cobalamin cofactor using Caver 3.0 (Chovancova *et al.* 2012). The models predicted that the I481T mutation is situated at the mouth of the major substrate access channel in each model (Figure 3B). Access channels can have a drastic impact on the substrate scope of an enzyme as they act as a filter for the active site. It has been established in directed evolution research that mutations in the substrate channel can lead to a change in enzyme activity (Kokkonen *et al.* 2019). Thus, it is speculated that the I481T mutation in TceA could alter substrate preferences.

The I481T mutation is also situated in a position where it can form interactions with one of the [4Fe-4S] clusters. While the level of influence that the [4Fe-4S] clusters and their redox potential have on RDase activity is unstudied, it is well known that the environment surrounding the clusters affects their redox potential (Langen *et al.* 1992; Birrell *et al.* 2016). The presence of hydrogen-bonding residues has been suggested to have an impact on the redox potential though there are other factors at play and a direct correlation with hydrogen bonding is not clear (Stephens, Jollie and Warshel 1996). The I481T mutation introduces the potential for a hydrogen bonding interaction and is predicted to be in close proximity (2.8 Å) of the [4Fe-4S] binding position (Figure 3C). As the redox potential provides a driving force for respiration and reduction of the electron acceptor, changes to the environment around the [4Fe-4S] clusters should be considered as a possible mechanism of fine-tuning the RDase activity towards a certain substrate.

Predictive modelling is a powerful tool in gaining insight as to the potential impact of these mutations; however, biochemical characterization and mutagenesis experiments need to be done to unequivocally determine the substrate range of the Donna II *D. mccartyi* strain 195 TceA. Furthermore, it would be beneficial to determine if mutations in TceA enhance VC respiration when combined with vitamin B_12_ supplementation, similarly to strain FL2 (Yan *et al.* 2021), or if TceA mutants can completely respire VC with lower concentrations of vitamin B_12_.

### Implications for horizontal gene transfer of the *vcrA*-GI

One objective of this experiment was to try to promote HGT of the *vcrA*-GI from *D. mccartyi* strain KBTCE1 into strain 195. As previously mentioned, HGT would be manifested by high/equal abundance of both KB-1/*vcrA* and 195/t*ceA* biomarkers with VC dichlorination (Table 1). These criteria were not met in any DKB sub-cultures. To further confirm that the *vcrA*-GI had not integrated into the *D. mccartyi* strain 195 genome, we PCR-amplified the integration site at the *ssrA* gene locus (McMurdie *et al.* 2011). We did not observe *vcrA*-GI integration in any DKB sub-cultures at the *ssrA* locus (Figure S7). As a result, we definitively concluded that HGT of the *vcrA*-GI did not occur in under the experimental conditions of the DKB experiment. In the family of KB-1 cultures enriched on different substrates, eight *D. mccartyi* genomes were closed (Molenda *et al.* 2020). These genomes revealed that clade-specific, and even strain-specific, characteristics provide *D. mccartyi* populations with ecological advantages to persist within mixed communities and to prevent gene transfer of mobile genetic elements between dissimilar strains. With the benefit of many more genome sequences, we now have evidence that *D. mccartyi* clade speciation plays a role in strain compatibility for HGT. To date, there are no Cornell strains that contain the *vcrA*-GI, only Pinellas and Victoria clades. There are several other clade-specific characteristics, including nitrogen fixation, which is found only in the Cornell clade, as previously described. HGT between clades may also be prevented by cellular defense mechanisms such as CRISPR-Cas systems (Molenda *et al.* 2019, 2020). With improved appreciation of the diversity of strains even within the KB-1/TCE-MeOH culture itself, there is also no evidence of *vcrA*-GI HGT between the Pinellas and Cornell strains now known to exist within this culture.

**Table 1:** Key features of *D. mccartyi* strains combined to try to promote horizontal gene transfer.

| Feature | *D. mccartyi* strain |  |  |
| --- | --- | --- | --- |
|  | KBTCE1<sup>a</sup> | 195 | Predicted “hybrid” |
| Role in HGT | Donor strain | Recipient strain | HGT product |
| Source Enrichment Culture | KB-1 | Donna II | DKB |
| Clade | Pinellas | Cornell | Cornell |
| *vcrA*-GI | *vcrA*-GI | -- | *vcrA*-GI |
| Nitrogen Fixation Operon | 0 | 1 | 1 |
| Distinguishing Biomarkers<sup>b</sup> | *vcrA* | *trdhA* | *vcrA* & *trdhA* |
| Common Biomarkers<sup>c</sup> | 16S | 16S, *tceA*, *nifD* | 16S, *tceA*, *nifD* |

<sup>a</sup>When the study was initiated, a *vcrA*-containing Pinellas strain was known to dominate the KB-1 TCE-enrichment culture. Subsequently, genomes corresponding to three strains (KBTCE1, KBTCE2, and KBTCE3) were closed from the KB-1/TCE-MeOH culture (Molenda *et al.* 2020). The KBTCE1 strain was most abundant when the culture was grown on TCE or VC.
<sup>b</sup>Key quantitative (q)PCR biomarkers used to distinguish strains.
<sup>c</sup>Biomarkers common to more than one strain, where 16S represents the *D. mccartyi* 16S rRNA gene.

### Starvation and long-term survival of *D. mccartyi* strains

These clade-specific and strain-specific characteristics may also play a role in the observed persistence of *D. mccartyi* strains and their long-term survival under starvation conditions. As previously mentioned, this finding may point toward RdhA of unknown function, such as the starvation RdhA (OG 15) which has not been functionally characterized but commonly observed in transcriptomic analyses (Johnson *et al.* 2008; Rahm and Richardson 2008; Waller *et al.* 2012; Tang *et al.* 2013). Furthermore, the long-term survival of *D. mccartyi* strains may explain why at least eight strains of *D. mccartyi* were identified in the KB-1 culture that has been maintained in the laboratory for more than 20 years (Molenda *et al.* 2020). The observations of the DKB experiment provide insights into the growth and survival of *D. mccartyi* in environments with low flow rates or where cells are attached to a surface, such as in the natural environment, that may also experience long-term starvation conditions. Organohalogens naturally exist in the environment, typically at low concentrations, and were the natural substrates for organohalide respiring bacteria before exposure to high concentrations of anthropogenic sources of organohalogens (Field 2016). It is likely that *D. mccartyi* have always been capable of long-term survival in the absence of organohalogens and these conditions have promoted strain variation among *D. mccartyi* communities.

While this experiment started off as a simple attempt to promote HGT, the breadth and depth of knowledge of *D. mccartyi* has developed immensely since the experiment began in 2012. In the time since the DKB culture was prepared at Cornell University, *D. mccartyi* was renamed (Löffler *et al.* 2013), the number of closed genomes of *D. mccartyi* in NCBI quintupled from 5 to 25 (Kube *et al.* 2005; Sung *et al.* 2006; Pöritz *et al.* 2013; Wang *et al.* 2014; Molenda, Quaile and Edwards 2016; Molenda *et al.* 2020), prophages (Waller *et al.* 2012), mobile genetic elements (Molenda *et al.* 2019), and CRISPR-Cas systems (Molenda *et al.* 2019) were identified in KB-1, and the genomes in the KB-1/TCE-MeOH (Molenda *et al.* 2020) and Donna II cultures (IMG-M taxon ID: 2032320001) were sequenced. Considering these advancements, the findings of the DKB experiment contribute to this knowledge and will inform attempts to induce HGT in the future.

## Acknowledgements

We acknowledge Garrett Debs, a researcher in the Richardson lab at Cornell University, who maintained the DKB in the early days, before it was sent to the University of Toronto.

