## Supplemental Information for "Long term survival of *Dehalococcoides mccartyi* strains in mixed cultures under electron acceptor and ammonium limitation"

### Supplemental Methods

#### Molecular Cloning for qPCR Standard Plasmids

For qPCR standards, primers were designed to capture regions up- and downstream from the qPCR target primers, using Primer3. After PCR amplification (PCR and primer details Table S2), PCR clean-up was performed using the QIAquick PCR Purification Kit (Qiagen). Following, PCR fragments were ligated into a cloning plasmid and transformed into *Escherichia coli* (Invitrogen TOPO® TA Cloning Kit with pCR®2.1-TOPO® vector and TOP10 cells). Plasmids from selected clones were extracted using the QIAprep Spin Miniprep Kit (Qiagen) and Sanger sequenced at the SickKids Centre for Applied Genomics TCAG sequencing facility (Toronto, Ontario) to confirm transformation. Subsequently, glycerol stocks of transformed E. coli were frozen (-80°C), and plasmids were extracted prior to qPCR analysis.

#### PCR strategy to track integration of the *vcrA*-GI at the *ssrA* locus

To observe when the *vcrA*-GI had integrated into a *D. mccartyi* genome, we designed a PCR reaction to amplify the insertion region of the *vcrA*-GI. The *vcrA*-GI is known to have site-specific integration downstream of *ssrA* in *D. mccartyi* genomes (1). This region also contains tandem genomic islands that are identifiable by 20 bp direct repeats of *ssrA* flanking the insertion. Additionally, integration at this site disrupts local gene synteny among *D. mccartyi* strains, and is identifiable by a change in %G+C content compared to the rest of the genome (1). However, some genomes of *D. mccartyi* do not contain any integrations at this site, for example strain 195. Therefore, a PCR reaction was designed to identify integrations at the *ssrA* integration locus. The PCR reaction was designed using the strain 195 genome as a template, where the forward primer targets the intragenic region between the *ssrA* gene (DET_RS07725) and the neighbouring gene (DET_RS07740). In the case of strain 195, when there are no integrations at the *ssrA* locus, the amplicon is 347 bp in length. However, these primers also amplify other *D. mccartyi* strains, which contain insertions at the *ssrA* locus, and in this case, depending on the integration, will vary in length. For example, when just the *vcrA*-GI is present, as in the case of the strain KBVC1 genome, originating from a VC-fed enriched KB-1 sub-culture (2), the amplicon is 11,692 bp in length.

The *ssrA* PCR reactions were performed in a 50 µL reaction using a high-fidelity polymerase (ThermoFisher Phusion, according to kit protocol) with the following program: initial denaturation at 98°C for 30 s, followed by 30 cycles of 10 s of denaturation at 98°C, annealing temperature 66°C, elongation at 72°C, and a final extension of 5 mins at 72°C. To analyze, the PCR products were separated on 1% TAE agarose gel, at 50 V for 1 hour, to estimate product size and check for non-specific amplifications. PCR products were sequenced (Sanger) at the SickKids Center for Applied Genomics (TCAG) sequencing/synthesis facility (Toronto, Ontario).

**Ammonium Quantification**

In DKB sub-cultures where ammonium was not provided in the medium, anaerobic mineral medium was prepared without the addition of $NH_4Cl$. To quantify the ammonium concentration in all experimental cultures throughout the experiment, ammonium was measured as $NH_3$-N by Nitrogen-Ammonia Reagent Set TNT AmVer Salicylate High Range kit (Hach), or as $NH_4^+$ by cation chromatography using the Dionex™ Integrion™ HPIC™ System (Thermo Fisher Scientific). For both preparations, 1 mL of each experimental culture was sampled anaerobically and aerobically injected into a 1.5 mL Eppendorf tube. For Hach kit quantification, a 100 µL aliquot of culture was diluted 1:3 in ammonium-free water and added to a kit test tube, the remainder of kit instructions were followed. A 10 mg/mL $NH_3$-N solution (Hach) was diluted to 1 mg/mL and 5 mg/mL for a 3-point calibration and used undiluted as an internal reference for each subsequent analysis. All kit test tubes were analyzed using the Hach DR3900 Spectrophotometer, with the built-in program #343 for absorbance at 655 nm. Final concentrations of $NH_3$-N were multiplied by the dilution factor and reported in mM concentrations. For cation chromatography, the 1 mL culture sample was filtered using a 0.2 µm pore size filter (13 mm diameter, Millex-GN Nylon membrane, hydrophilic, Millipore Sigma). A 500 µL aliquot was pipetted into 0.5 mL Dionex™ AS-DV Autosampler PolyVials (Thermo Fisher Scientific). Method calibration was established using a 5-point standard curve of 0.005-1 mM concentrations of $NH_4Cl$, using a CS19 column, with an isocratic method of 8 mM methane sulfonic acid as eluent, flow rate of 1mL/min, and suppressor voltage of 3.9 V. The line of best fit of the standard curve was a polynomial, 2$^{nd}$ degree line ($y = 0.101x^2 + 0.164x$). $NH_4^+$ had a retention time between 4.5 and 4.6 minutes.

### Supplemental Figures

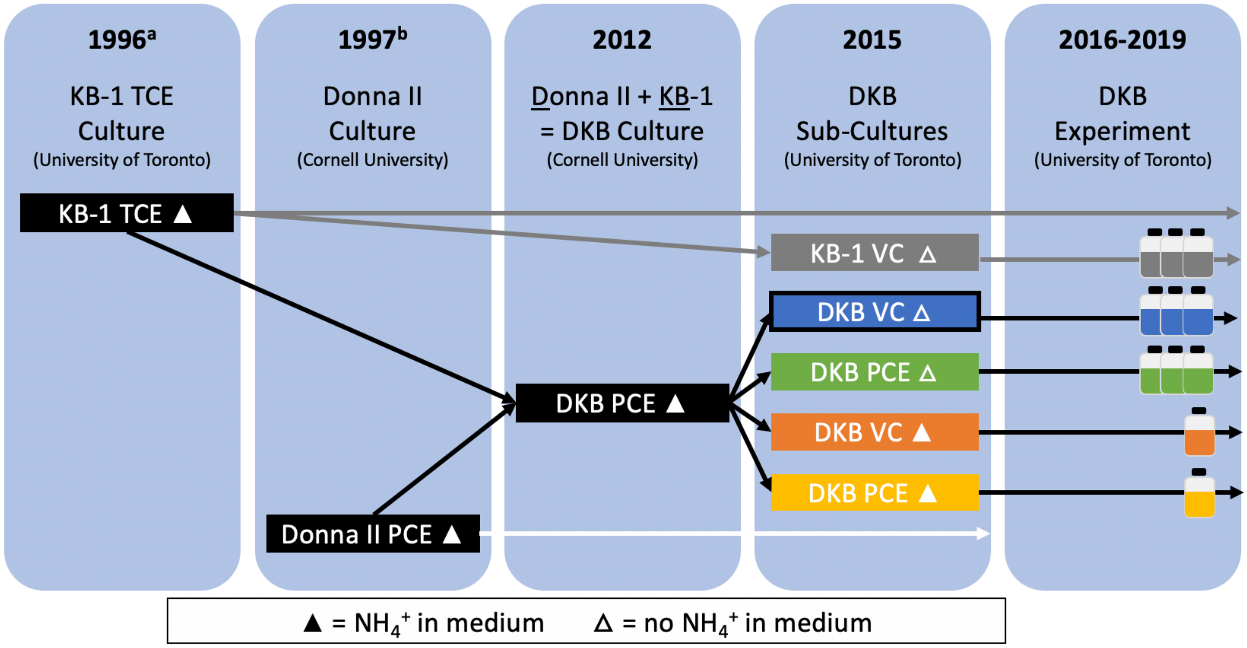

**Figure S1:** DKB experiment timeline. The KB-1 TCE-enriched culture [a](3) and the Donna II culture [b](4) were combined to create the DKB culture in 2012. KB-1 sub-cultures ("KB-1 VC △", grey) were created as a negative control for nitrogen fixation. The DKB sub-culture condition with the highest selective pressure was provided VC as the sole energy source without providing ammonium ("DKB VC △", blue box with black border), was predicted to be the most likely condition to promote HGT of the *vcrA*-GI. The number of bottles in the last panel indicate how many sub-culture replicates were maintained and analyzed during this experiment (2016-2019).

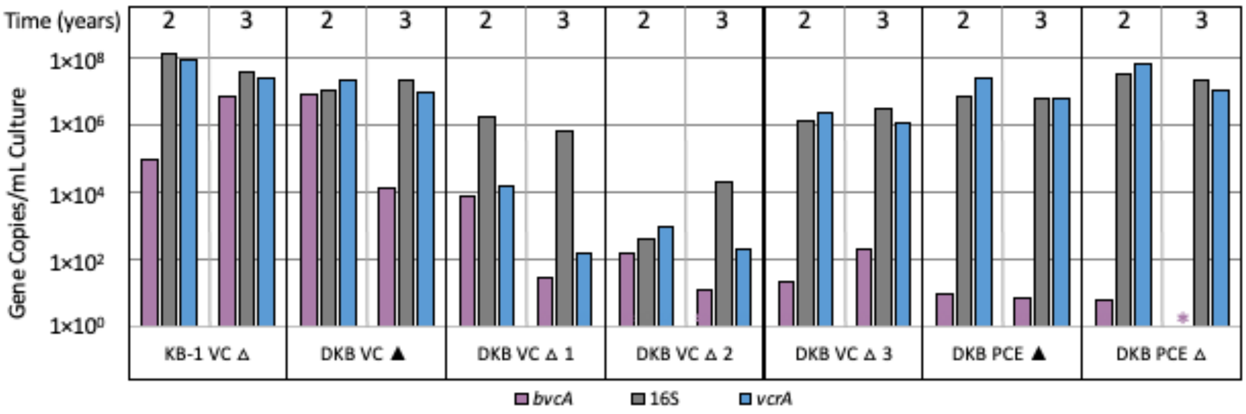

**Figure S2:** Quantification of *bvcA* relative to *D. mccartyi* 16S rRNA, and *vcrA* in the KB-1 VC ∆ cultures and DKB sub-cultures. The colored * indicates the gene was not detected. The quantification limit was $1\times10^3$ copies/mL; values below this are unreliable. Where ▲ indicates the culture was grown with ammonium provided and ∆ indicates the culture was grown without ammonium provided. The number indicates the replicate bottle number.

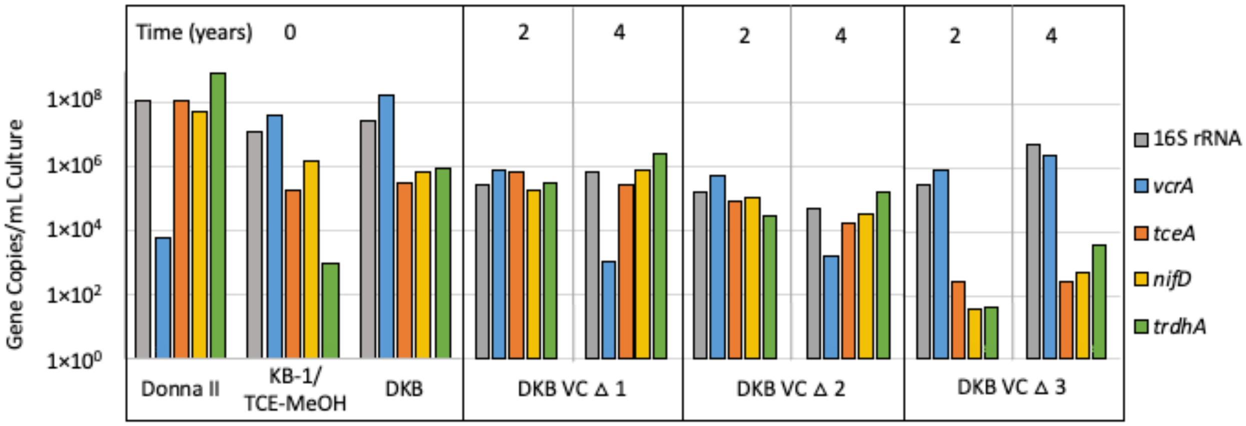

**Figure S3:** *D. mccartyi* strain-specific biomarkers in the DKB VC ∆ cultures. Donna II, KB-1/TCE-MeOH and DKB cultures at time 0 and DKB VC ∆ cultures during years 2 and 4. The quantification limit was $1\times10^3$ copies/mL; values below this are unreliable. Where ▲ indicates the culture was grown with ammonium provided and ∆ indicates the culture was grown without ammonium provided. The number indicates the replicate bottle number.

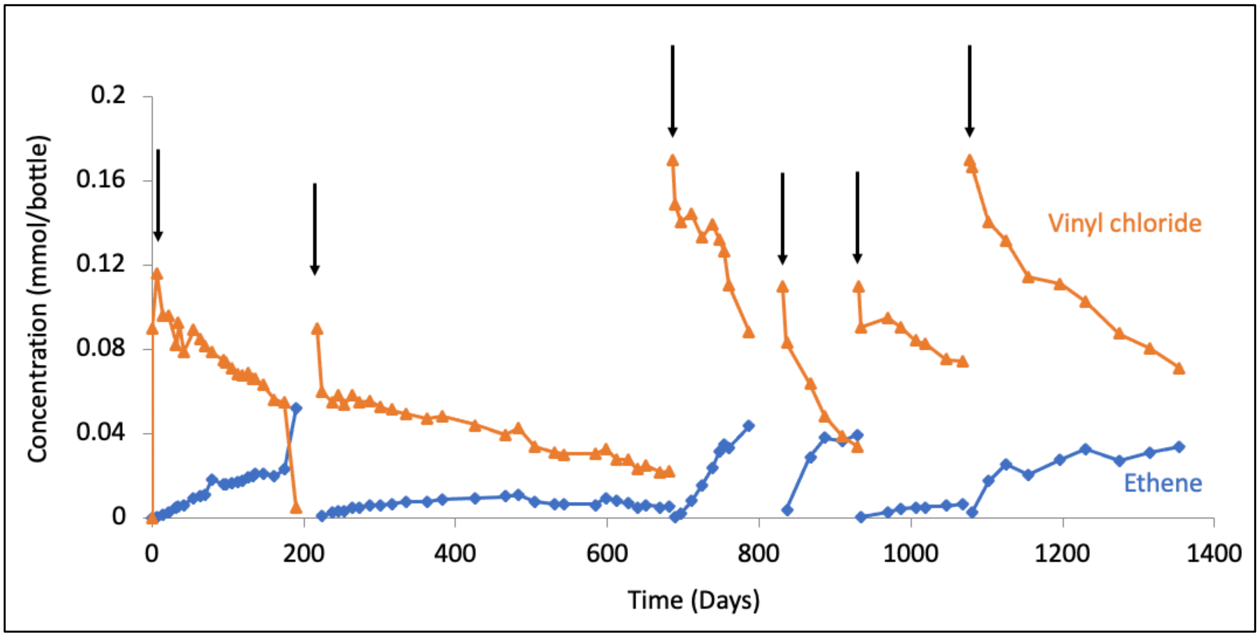

**Figure S4:** Vinyl chloride dechlorination to ethene in the DKB VC ∆ 1 culture, measured by gas chromatography. The VC dechlorination rate for the last dechlorination cycle (days 1081 and 1354) was 1.4 µmol VC dechlorinated/L/day. Black arrows indicate where VC-feeding occurred.

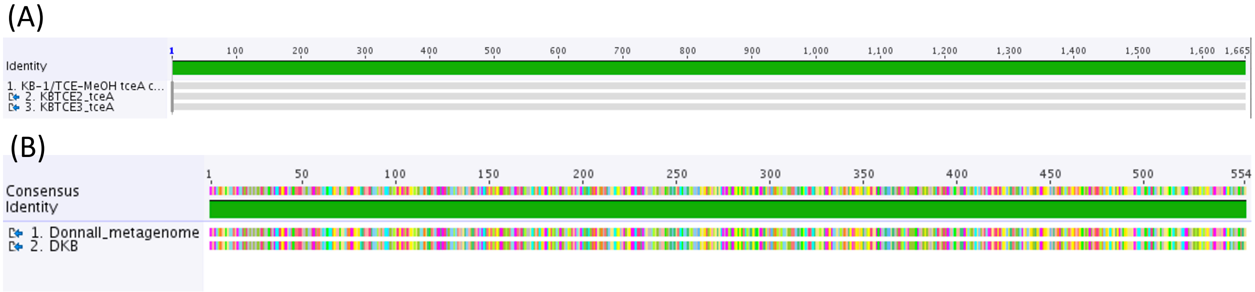

**Figure S5:** Cloned *tceA* aligned to parent culture *tceA* sequences. (A) KB-1/TCE-MeOH *tceA* clone and *D. mccartyi* strains KBTCE2 and KBTCE3 tceA alignment. The green identity bar represents 100% nucleotide pairwise identity. (B) Donna II metagenome TceA and DKB VC ∆ 1 TceA clone. The green identity bar represents 100% amino acid pairwise identity.

A

| | 259 | 260 | 261 | 262 | 263 |
|---|---|---|---|---|---|
| 195 | A | K | V | Q | P |
| Donna II | A | K | A | Q | P |
| DKB | A | K | A | Q | P |
| KBTCE2 | T | K | A | Q | P |
| KBTCE3 | T | K | A | Q | P |
| FL2 | A | K | A | Q | P |
| UCH-ATV1 | T | K | A | Q | P |
| BTF08 | T | K | A | Q | P |
| 11a5 | T | K | A | Q | P |

B

| | 479 | 480 | 481 | 482 | 483 |
|---|---|---|---|---|---|
| 195 | K | C | I | N | C |
| Donna II | K | C | T | N | C |
| DKB | K | C | T | N | C |
| KBTCE2 | K | C | I | N | C |
| KBTCE3 | K | C | I | N | C |
| FL2 | K | C | T | N | C |
| UCH-ATV1 | K | C | I | N | C |
| BTF08 | K | C | I | N | C |
| 11a5 | K | C | I | N | C |

**Figure S6:** Protein alignment of TceA compared to the *D. mccartyi* strain 195 isolate sequence. Alignment includes all TceA in OG 5 (5), with the addition of TceA from the Donna II metagenome (IMG-M ID: 2032320001) and the cloned TceA from the DKB experiment (DKB VC ∆ 1). (A) TceA SNP at position 261, where there is a Val (orange) in the strain 195 isolate sequence, but an Ala (blue) in all other TceA sequences. (B) TceA SNP at position 481

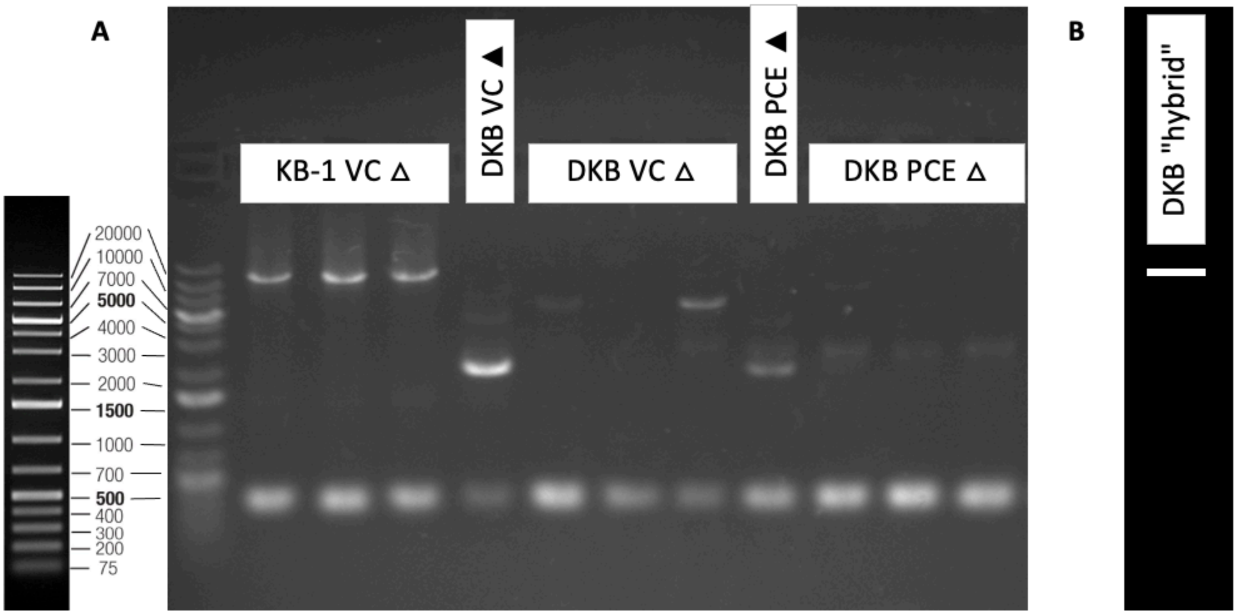

**Figure S7:** PCR to determine integration of the *vcrA*-GI at the *ssrA* locus in KB-1 and DKB cultures. Thermo Scientific™ GeneRuler™ 1 kb Plus DNA Ladder (left) on 1% agarose gel. (A) PCR amplicons were ~11 kbp when the *vcrA*-GI was integrated into the genome, and 346 bp when it was not. All amplicons between 11 kb and 346 bp were cloned, sequenced and identified as non-specific PCR amplification. The integration of the *vcrA*-GI at the *ssrA* locus was only found in the KB-1 VC ∆ replicates. (B) The DKB "hybrid" strain, as a result of the HGT event, would have amplified the 11 kbp PCR product, which would have indicated that the *vcrA*-GI transferred into the *D. mccartyi* strain 195 genome. This amplicon was not identified in any DKB sub-cultures.

**Supplemental Tables**

Table S1 [excel]: Cumulative eeq consumed in DKB experimental bottles, based on calculated mass of electron acceptor fed and time to consume.

Table S2 [excel]: Quantitative (q)PCR and cloning primers used for molecular analysis.

Table S3 [excel]: Quantitative (q)PCR standard curves generated for quantification of biomarker genes.

Table S4 [excel]: Ammonium concentration in DKB experiment using a Hach Nitrogen-Ammonia kit.

Table S5 [excel]: Absolute concentrations of biomarkers by quantitative (q)PCR analyses.

Table S6 [excel]: Nitrogenase operons of *D. mccartyi* Cornell strains KBTCE2, KBTCE3 and 195.

Table S7 [excel]: Comparison of *nifD* transcripts and gene copies per mL of KB-1, 195 and PW4 cultures and TPG calculations.

Table S8 [excel]: Complete nucleotide and protein sequences of the Donna II metagenome strain 195 TCE reductase.

Table S9 [excel]: Quality scores of isolate strain 195 and Donna II strain 195 TceA models using Phyre2, I-TASSER and Robetta.
